# Spontaneous isomerization of long-lived proteins provides a molecular mechanism for the lysosomal failure observed in Alzheimer’s disease

**DOI:** 10.1101/605626

**Authors:** Tyler R. Lambeth, Dylan L. Riggs, Lance E. Talbert, Jin Tang, Emily Coburn, Amrik S. Kang, Jessica Noll, Catherine Augello, Byron D. Ford, Ryan R. Julian

## Abstract

Proteinaceous aggregation is a well-known observable in Alzheimer’s disease (AD), but failure and storage of lysosomal bodies within neurons is equally ubiquitous and actually precedes bulk accumulation of extracellular amyloid plaque. In fact, AD shares many similarities with certain lysosomal storage disorders though establishing a biochemical connection has proven difficult. Herein, we demonstrate that isomerization and epimerization, which are spontaneous chemical modifications that occur in long-lived proteins, prevent digestion by the proteases in the lysosome (namely the cathepsins). For example, isomerization of aspartic acid into L-isoAsp prevents digestion of the N-terminal portion of Aβ by cathepsin L, one of the most aggressive lysosomal proteases. Similar results were obtained after examination of various target peptides with a full series of cathepsins, including endo-, amino-, and carboxy-peptidases. In all cases peptide fragments too long for transporter recognition or release from the lysosome persisted after treatment, providing a mechanism for eventual lysosomal storage and bridging the gap between AD and lysosomal storage disorders. Additional experiments with microglial cells confirmed that isomerization disrupts proteolysis in active lysosomes. These results are easily rationalized in terms of protease active sites, which are engineered to precisely orient the peptide backbone and cannot accommodate the backbone shift caused by isoaspartic acid or side chain dislocation resulting from epimerization. Although Aβ is known to be isomerized and epimerized in plaques present in AD brains, we further establish that the rates of modification for aspartic acid in positions 1 and 7 are fast and could accrue prior to plaque formation. Spontaneous chemistry can therefore provide modified substrates capable of inducing gradual lysosomal failure, which may play an important role in the cascade of events leading to the disrupted proteostasis, amyloid formation, and tauopathies associated with AD.

## Introduction

The active balancing of protein synthesis and degradation, or proteostasis, is an ongoing and critical process in most cells.^1^ Proteins must be created, carry out their requisite function, and then be recycled once they are no longer needed or have become nonfunctional. Several pathways are available for protein degradation, including the proteasome, macro-autophagy, micro-autophagy, and chaperone-mediated autophagy.^2,3^ The autophagy-related pathways deliver proteins to lysosomes, which are acidic organelles containing a host of hydrolases, including many proteases.^4^ Cargo taken into cells via endocytosis is also typically delivered to lysosomes for degradation. Regardless of the pathway, after cargo fuses with a lysosome, endopeptidases cleave proteins at internal sites, shortening proteins to peptides, which are then further digested from both termini by exopeptidases. After protein digestion has been completed, transporter proteins in the lysosomal membrane release (primarily) individual amino acids back into the cytosol for new protein synthesis or energy production.^5^ Lysosomes are crucial for maintaining cellular homeostasis, but they are also uniquely susceptible to problems when substrates cannot be hydrolyzed. For example, genetic modifications reducing the efficacy of a lysosomal hydrolase are the most common cause of lysosomal storage disorders. These devastating diseases involve ‘storage’ of failed lysosomal bodies within cells, which eventually leads to cell death and is particularly problematic for post-mitotic cells such as neurons.^6^ Symptoms in lysosomal storage disorders usually emerge in infancy or childhood, are often associated with neurodegeneration, and are typically fatal.^7^

Long-lived proteins^8^ are a primary target of the lysosome because they become modified and lose efficacy over time. A well-known example of this occurs with mitophagy,^3^ wherein old mitochondria are recycled in their entirety. Contributing factors that lead to long-lived protein deterioration include a variety of spontaneous chemical modifications, i.e. modifications not under enzymatic control.^8^ Some of these modifications are very subtle and difficult to detect, including isomerization and epimerization.^9^ Isomerization occurs primarily at aspartic acid, when the side chain inserts into and elongates the peptide backbone (Scheme S1).^10^ Identical products are also created during deamidation of asparagine, which further results in chemical transformation from one amino acid to another.^11^ Epimerization occurs when an amino acid side chain inverts chirality from the L- to D-configuration. Peptide isomerization and epimerization do not have readily identifiable bioanalytical signatures, but both modulate structure in a subtle, yet significant way (see Fig. 1). Studies on the eye lens have shown that epimerization and isomerization are among the most abundant modifications observed in extremely long-lived proteins.^12–14^ However, knock-out experiments in mice have also revealed the importance of these modifications over much shorter timescales. For example, removal of the repair enzyme for L-isoAsp, protein-isoaspartyl methyl transferase (PIMT),^15^ leads to lethal accumulation of isomerized protein in just 4-6 weeks.^16,17^ This reveals that isomerization of aspartic acid is sufficiently dangerous that an enzyme has evolved to repair it.

**Fig.1.**
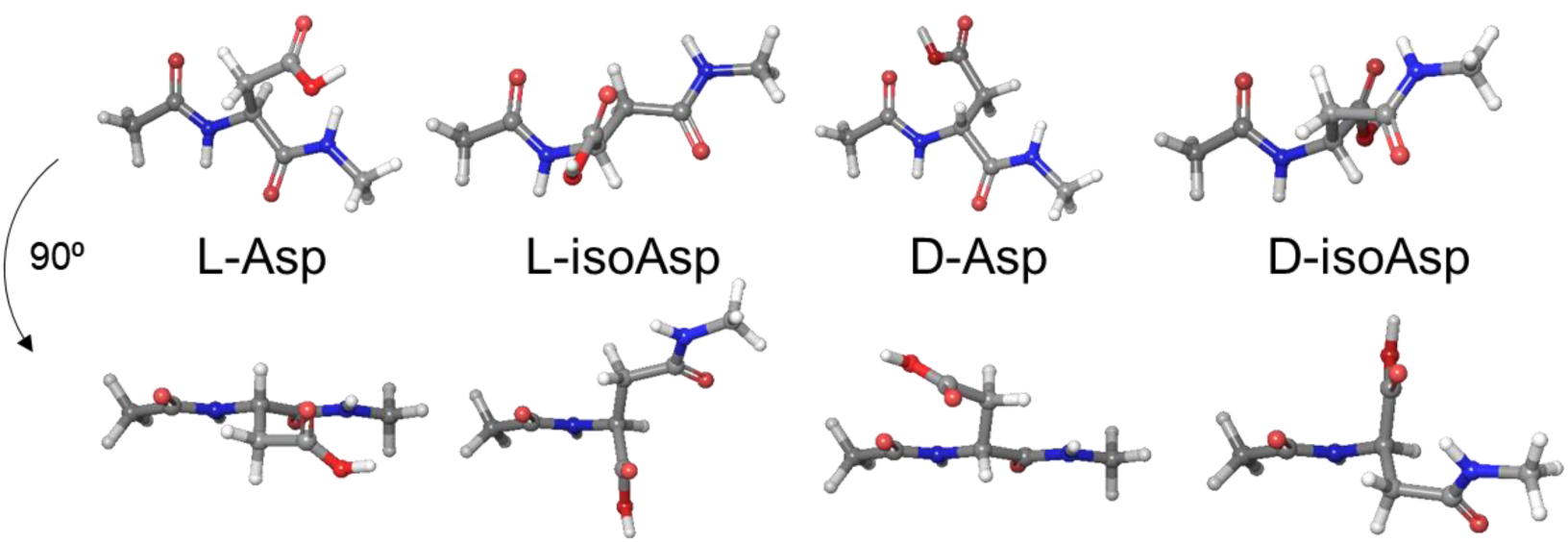
Model structures of the aspartic acid isomers, where the iso-structure conformation closest to native backbone orientation is shown. Two views are illustrated for each isomer.

The importance of peptide isomers is further revealed in the uses nature has found for them. For example, single amino acid sites are intentionally epimerized in many venoms and in signaling neuropeptides in crustaceans.^18,19^ The corresponding L-only peptides are not biologically active, confirming the importance of the chiral modifications. In addition, it is thought that epimerization is beneficial for these peptides because it allows them to escape or prolong the time required for proteolysis.^20^ In fact, it is well-known that sites of epimerization and isomerization are both generally resistant to protease action, but the ramifications of such chemistry in the context of lysosome function have not been previously examined. Despite this absence, numerous studies have the established the importance of protein degradation in lysosomes. For example, knockout mice lacking cathepsin D grow normally for ~2 weeks but then die before the end of 4 weeks.^21^ Examination of the neurons from these mice revealed an abundance of failed lysosomal bodies, similar to those observed in lysosomal storage disorders. Other research has shown that knockout mice lacking cathepsins B and L die within 2-4 weeks of birth. Again, accumulation of failed lysosomal bodies was observed in neurons of these mice.^22^ Although cathepsins can also be found outside the lysosome, these results confirm significant, and likely fatal, impact on the lysosomal system when critical cathepsins are absent.

Amyloid aggregates or proteins that are otherwise insoluble are also targeted to lysosomes for degradation. Amyloid aggregation has also captured the majority of attention as the potential cause of Alzheimer’s disease (AD), but significant evidence also supports lysosomal storage as an underlying cause. For example, AD shares many pathological similarities with lysosomal storage disorders, including prolific storage of failed lysosomal bodies, accumulation of senile plaques, and formation of neurofibrillary tangles.^23,24^ In fact, scanning-electron microscopy images of lysosomal storage (in neurons) are virtually indistinguishable between the two diseases. The lysosomal storage observed in AD precedes formation of amyloid deposits,^25^ hinting that lysosomal malfunction may occur upstream of the events leading to amyloid aggregation. The parallels between the two diseases have also been offset by differences. For example, lysosomal storage disorders typically afflict youth and can progress rapidly, while AD typically occurs late in life over a longer timescale. Therefore, a mechanism accounting for the commonalities and differences between the diseases has been difficult to identify, but an intriguing possibility does exist.

The primary constituents of senile plaques, Aβ and tau, are both long-lived proteins that are subject to isomerization and epimerization. In fact, Aβ is significantly epimerized and isomerized in the brains of people with AD.^26^ If isomerization and epimerization prevent lysosomal protein digestion, then a common link between lysosomal storage disorders and AD would be established. In fact, AD would essentially represent a type of lysosomal storage disorder, one that operates in reverse of the classical disease. Rather than failure of a *modified enzyme or modified transporter* to clear waste molecules, failure to digest or transport *modified waste* molecules would be operative and eventually lead to lysosomal storage. Close examination of another complex age-related disease, macular degeneration, reveals that there is precedence for substrate-induced lysosomal storage.^27^

Herein, we use mass spectrometry (MS) and liquid chromatography (LC) to demonstrate that isomerized or epimerized peptides resist degradation by cathepsins, including both endo- and exopeptidase activity. Important target peptides that are both long-lived and closely associated with AD were examined, including fragments of Aβ and tau. The results reveal that small peptide fragments comprised of residues surrounding isomerized or epimerized sites will persist after digestion. Disrupted proteolysis was observed in both isolated reactions and living cells and offers an explanation for toxicity observed in previous experiments with cell and animal models employing isomerized Aβ. Additional experiments reveal that the rates of isomerization for the N-terminal Asp residues in Aβ are fast, providing a pathway for generation of these toxic species that could eventually lead to lysosomal failure and initiate other downstream consequences.

## Results and Discussion

### Defining limitations of cathepsin digestion

A series of isolated digestions of synthetic peptides in both canonical form and with isomerized or epimerized (iso/epi) sites were performed and the results are summarized in Fig. 2. Experiments were conducted with cathepsins D, L, B, and H. This collection includes all of the most abundant cathepsins and all modes of function, i.e. endo-, carboxy-, and aminopeptidases.^28,29^ The peptide APSWFDTGLSEMR (αB^57–69^), derived from αB-crystallin, was used as the initial test substrate. It contains both Ser and Asp residues known to be modified in the eye lens.^30^ Furthermore, each iso/epi site is isolated, allowing for semi-independent examination, and the canonical sequence is a good substrate for proteolysis. Digestion of the native form with cathepsin D yields the results shown in the upper part of Fig. 2a. The LC-MS derived ion chromatogram reveals many peptide fragments and almost complete consumption of the precursor. Clearly, the canonical all-L version of APSWFDTGLSEMR is easily digested. Substitution of L-Asp with L-isoAsp yields the lower chromatogram, where after 6 hours, the precursor remains basically untouched. A single modification therefore prevents cathepsin D from digesting an entire thirteen residue sequence, shutting down peptide hydrolysis at seven different sites. To better visualize the results, peptide fragments resulting from proteolysis are represented by color-coded lines below the peptide sequence as shown in Fig. 2b. The data from Fig. 2a corresponds to the top two rows of the results shown in Fig. 2b. Data for the other Asp isomers and both Ser epimers are shown in the remaining slots of Fig. 2b. All three non-native forms of aspartic acid essentially prevent digestion by cathepsin D. Furthermore, epimerization of the less bulky serine side chain also modulates cathepsin D action, preventing cleavage at one or more preferred sites even when the epimerized serine is located six residues away. Significant residual precursor is detected for all modifications, suggesting decreased affinity for the iso/epi modified peptides in general. Results for analogous experiments with cathepsin L are shown in Fig. 2c. The canonical peptide is digested into many peptide fragments, including small di- and tripeptides. Cathepsin L is one of the most aggressive lysosomal proteases and is able to cleave more sites in the iso/epi modified peptides relative to cathepsin D. Furthermore, precursor survival is not observed with cathepsin L. However, the sites where digestion occurs are all shifted well away from iso/epi modified residues in every instance, and the number of peptide fragments observed is still reduced relative to the canonical form. The results from cathepsin L and D reveal that digestion by endopeptidase action is significantly hampered by iso/epi modifications across wide regions of sequence.

**Figure 2.**
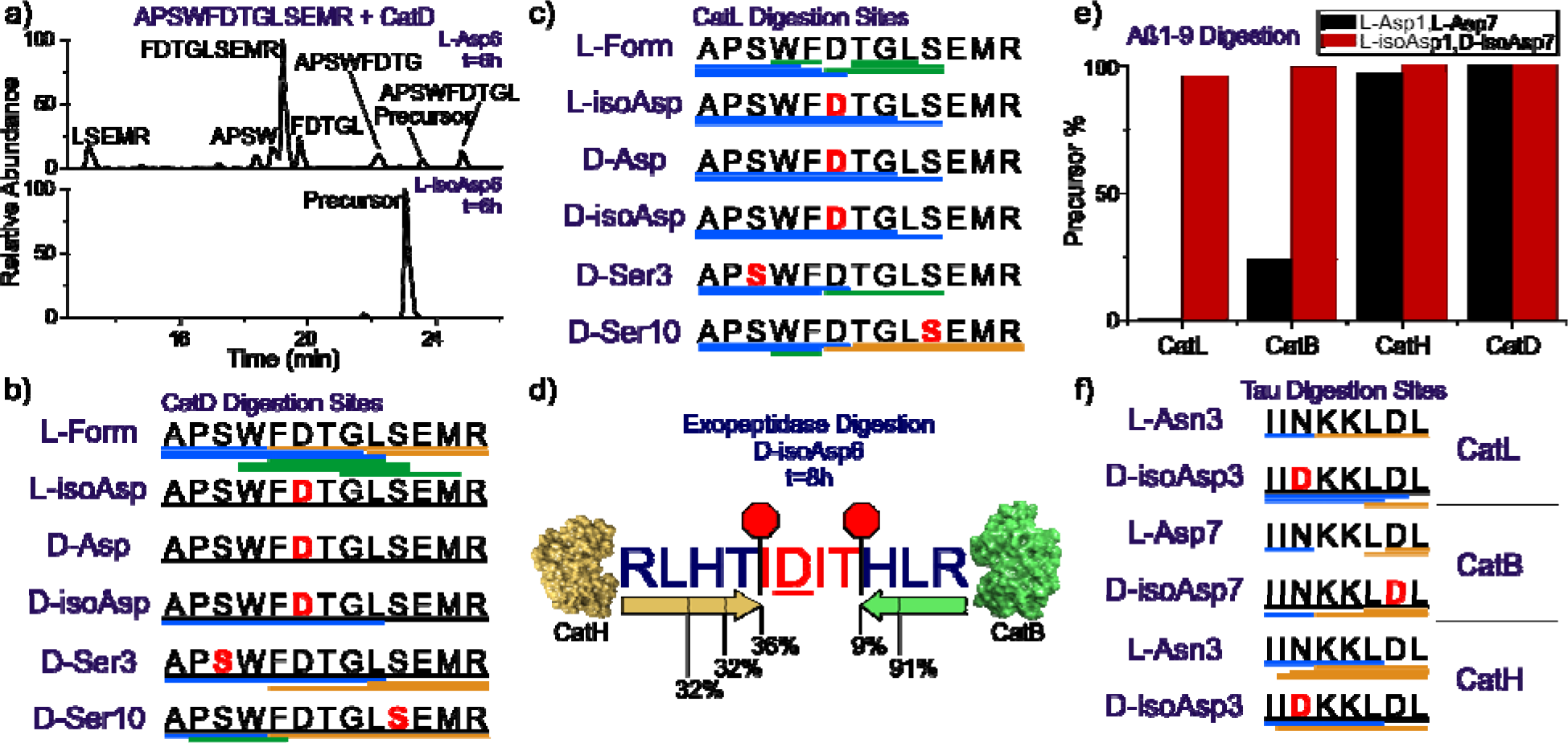
a) LC chromatogram for digestion of APSWFDTGLSEMR by cathepsin D. Summary of digestion by b) cathepsin D and c) cathepsin L. Each bar represents a fragment detected in the LC-MS chromatogram, color coded by N-terminal (blue), C-terminal (gold), and internal (green). Undigested precursor >50% relative intensity is represented by a black line. d) Summary of digestion by exopeptidases cathepsin H and B, illustrating residual core. e) Summary of digestion of Aβ1-9 (L-Asp1, L-Asp7) vs (L-isoAsp1, D-isoAsp7) by major cathepsins. Only the canonical isomer is digested. f) Summary of digestion of ^594^IINKKLDL^601^ from Tau using same color scheme.

The lysosomal task of reducing proteins and peptides into individual amino acids is never completed by endopeptidases, making examination of exopeptidases important. We used a palindromic peptide (RLHTIDITHLR) to systematically explore the limits of exopeptidase activity, and the results for experiments with cathepsins H and B are summarized schematically in Fig. 2d. Although both exopeptidases are able to advance in from either termini, the IDIT fragment persists even after 48 hours digestion time. As an exopeptidase, cathepsin B preferentially removes dipeptides, while cathepsin H behaves as an aminopeptidase, removing a single N-terminal amino acid at a time.^28^ It is not surprising that cathepsin H is able to penetrate closer to the iso/epi site because it does not need to accommodate two amino acids into the catalytic site. Taken together, the results illustrate significant disruption of proteolysis by iso/epi modifications for both the major endo- and exo-lysosomal peptidases.

The results for additional peptide targets relevant to AD are shown in Fig. 2e and 2f (Aβ1-9 and Tau ^594^IINKKLDL^601^). The two aspartic acids near the N-terminus of Aβ, Asp1 and Asp7, are highly isomerized in amyloid plaques,^26^ and represent an interesting target where multiple proximal iso/epi modifications can be found. Isomerization of Aβ is known to inhibit serum protease action, suggesting that cathepsins may likewise be stymied.^31^ Experiments conducted on canonical Aβ1-9 and a double isomer (L-isoAsp1, D-isoAsp7) are summarized Fig. 2e, where the fraction of remaining precursor from each peptide is shown for each cathepsin. Cathepsin B and L easily deplete the precursor for the canonical peptide, but are unable to significantly reduce the amount of precursor for the double isomer. Interestingly, cathepsin D cleaves few sites^32^ in Aβ and is unable to cleave any portion of Aβ1-9 even in canonical form. Similarly, cathepsin H exhibits low affinity for the N-terminal residues in Aβ1-9, and digests the canonical peptide only marginally while leaving the isomerized form intact. The N-terminal portion of Aβ is therefore generally resistant to lysosomal protease action and home to multiple sites of modification that can further frustrate proteolysis, making the prospects for Aβ to contribute to lysosomal failure strong.

Tau-mediated pathology is also strongly associated with AD, making it an important target to consider.^33^ Asn596 in Tau is known to deamidate,^34^ which will yield conversion to Asp and iso/epi modifications according the pathway illustrated in Scheme S1. As a long-lived protein, Tau could also isomerize at Asp600. Isomerization at both sites is explored for the peptide fragment ^594^IINKKLDL^601^ in Fig. 2f for cathepsins B, L, and H. The canonical peptide is rapidly consumed for all three cathepsins, but introduction of D-isoAsp at either position significantly perturbs the locations of proteolytic cleavage sites and leads to observation of abundant undigested precursor in all cases. These results reveal that inhibited proteolysis is likely a general feature of iso/epi modified peptides, and sequences known to be modified in the brain will be difficult to digest in the lysosomal pathway.

### Isomer digestion in living cells

To explore additional lysosomal proteases, experiments were conducted with fully active lysosomes in SIM-A9 mouse microglial cells, as shown in Fig. 3. For the peptide target, the N-terminal portion of Aβ was selected, and microglial cells were used because they are active participants in the clearance of Aβ within the brain.^35^ Chimeric peptides (R_8_-*E*_*edan*_DAEFRHD*K*_*dab*_G, where the Glu and Lys have been modified with edans and dabcyl, respectively) consisting of a cell-penetrating portion combined with an Aβ probe sequence were synthesized. Polyarginine was used for cell penetration, which is known to deliver cargo to the lysosome.^36^ The probe portion of the peptide remains dark when intact as the edans fluorescence is efficiently quenched by dabcyl. Upon cleavage of the probe sequence, the quencher can separate and edans will emit broadly around 490nm. Results for Aβ1-7 (L-Asp1, L-Asp7) as the probe are shown in Fig. 3a, revealing that fluorescence is observed after 150 min as expected. In comparison, the D-isoAsp1/D-isoAsp7 probe yields lower intensity fluorescence in terms of quartile range, median, and number (including exceptionally bright cells), as shown in Fig. 3b. Statistical comparison of the results with the Mann-Whitney U test reveals that differences in digestion are significant for all time points. These results suggest that there is not an unknown protease in the lysosome engineered to digest iso/epi sites.

**Figure 3.**
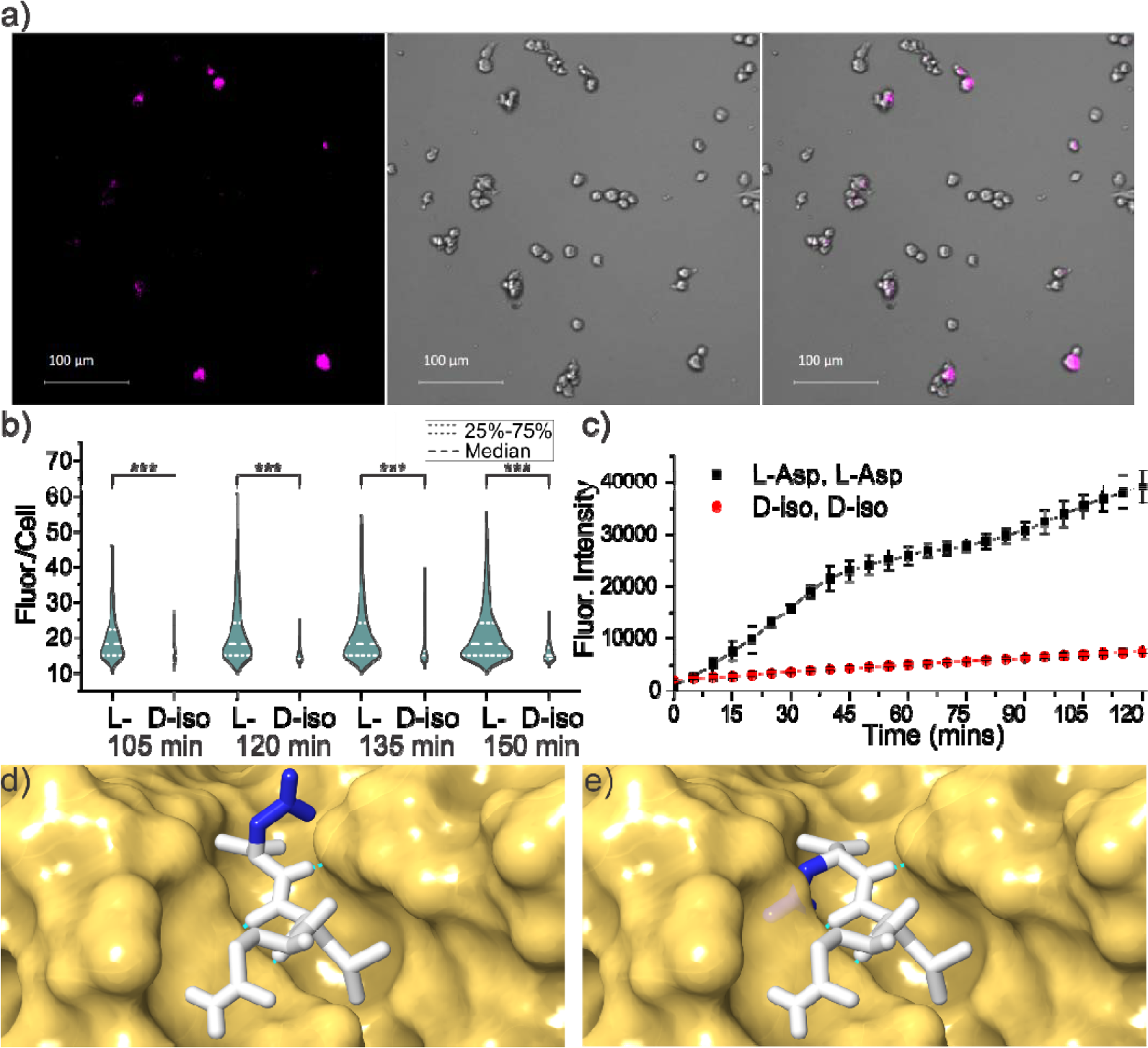
a) Sample images of SIM-A9 mouse microglial cells after 150 minute incubation with cleavable peptide target with all L-residues, fluorescence from 481-499 nm (left), brightfield (middle), and overlay (right). b) Violin plot showing quantitative comparison of fluorescence intensity per cell from Aβ1-7 cleavage for canonical and the D-isoAsp1/D-isoAsp7 isomers as a function of incubation time. *** p < 0.001 c) Fluorescence intensity as a function of time for incubation of same peptide with cathepsin L only. d) Active site of cathepsin L with native peptide substrate bound and e) mutated epimer with D-Asp sidechain highlighting inherent steric clash if backbone orientation is maintained. Structures derived from PDB ID: 3K24 with hydrogen bonds indicated by green dashed lines.

Interestingly, the microglial results can be largely recapitulated by examination of the same chimeric peptide incubated with only cathepsin L, as shown in Fig. 3c and Fig. S1. Both the rates and magnitude of the differential closely match the results obtained in living cells. These findings are consistent with previous observations that cathepsin L is one of the most important lysosomal proteases and can account for ~40% of all protein digestion in the lysosome.^28^ The accurate reproduction confirms the validity of the LC-MS approach that yielded the results shown Fig. 2. Furthermore, the effects of iso/epi modifications are more accurately determined under controlled incubation where canonical peptides without additional modifications can be tested. For example, Aβ1-7 (D-isoAsp, D-isoAsp) itself is almost completely resistant to degradation, yet proteolysis with cathepsin L is increased by a factor of ~7 after decoration with hydrophobic chromophores needed for examination in cells (Fig. S2). This suggests that the difference between digestions shown in Fig. 3b is significantly underestimated relative to the true inhibiting power of the D-isoAsp modifications.

The results in Fig. 2 and 3a,b can easily be rationalized by a molecular level inspection of the interaction between a protease and substrate peptide. In Fig. 3d, the X-ray crystal structure for binding of a peptide substrate to cathepsin L is shown.^37^ The protease active site consists of a channel where several hydrogen bonds orient the peptide backbone of the substrate. Intimate contact and alignment of the substrate backbone is required to bring the cleavage site into proximity with the catalytic actors. Favorable or unfavorable interactions with side chains protruding above the groove determine the sequence selectivity, but introduction of a D-amino acid with the peptide backbone remaining properly oriented would result in the side chain projecting directly into the wall of the binding groove (Fig. 3e). Similarly, isoAsp modifications disrupt both the backbone hydrogen bond partner spacing and relative orientation (Fig. 1), making for an even less tractable situation. These structural alterations make it impossible for iso/epi modified residues to fit properly into the catalytic binding site. Given the similarities inherent in the function and substrate for every protease, comparable complications are likely to exist for all proteases intended to cleave peptides comprised solely of canonical L-residues. Perhaps it is not surprising that poor proteolysis is observed for iso/epi modified peptides even in glial cells where a full complement of lysosomal proteases is available.

### Timeframe for aspartic acid isomerization

Given that Aβ plays an important role in Alzheimer’s disease (AD) and is highly isomerized in amyloid plaques,^26,38^ we set out to determine the incubation times needed to yield such extensive modifications. Following incubation of Aβ 1-40, 1-42, and 1-9 in tris buffer at 37ºC, the degree of isomerization is shown as a function of time in Figs. 4a and 4b. The isomerization of Asp1 and Asp7 were determined independently by digesting the aged products with chymotrypsin, yielding ^1^DAEF^4^ and ^5^RHDSGY^10^ peptides. Isomerization occurs rapidly at both aspartic acids for both full length peptides, yielding roughly 14% combined isomerization within 30 days. This rate is comparable to previous examination^39^ of Aβ1-16 and to isomerization of Asp151 in αA crystallin (when determined for the peptide fragment ^146^IQTGLDATHAER^157^). ^40^ It is also consistent with other isomerization rates cited in the literature as shown in Fig. 4c,^41–44^ where the only significantly faster rates involve Asp-Gly sequences. Detailed study of deamidation, which forms an identical succinimide ring intermediate preceding isomerization, revealed the fastest rates for analogous Asn-Gly sites.^45^

^1^Ref 41

^2^Ref 40

^3^Ref 42

^3b,c^Estimated rate of the VYPDGA peptide from Literature point 3 modified to correspond to VYPDSA and VYPDAA based on known deamidation rates.^45^

^4^Ref 43

^5^Ref 44

**Figure 4.**
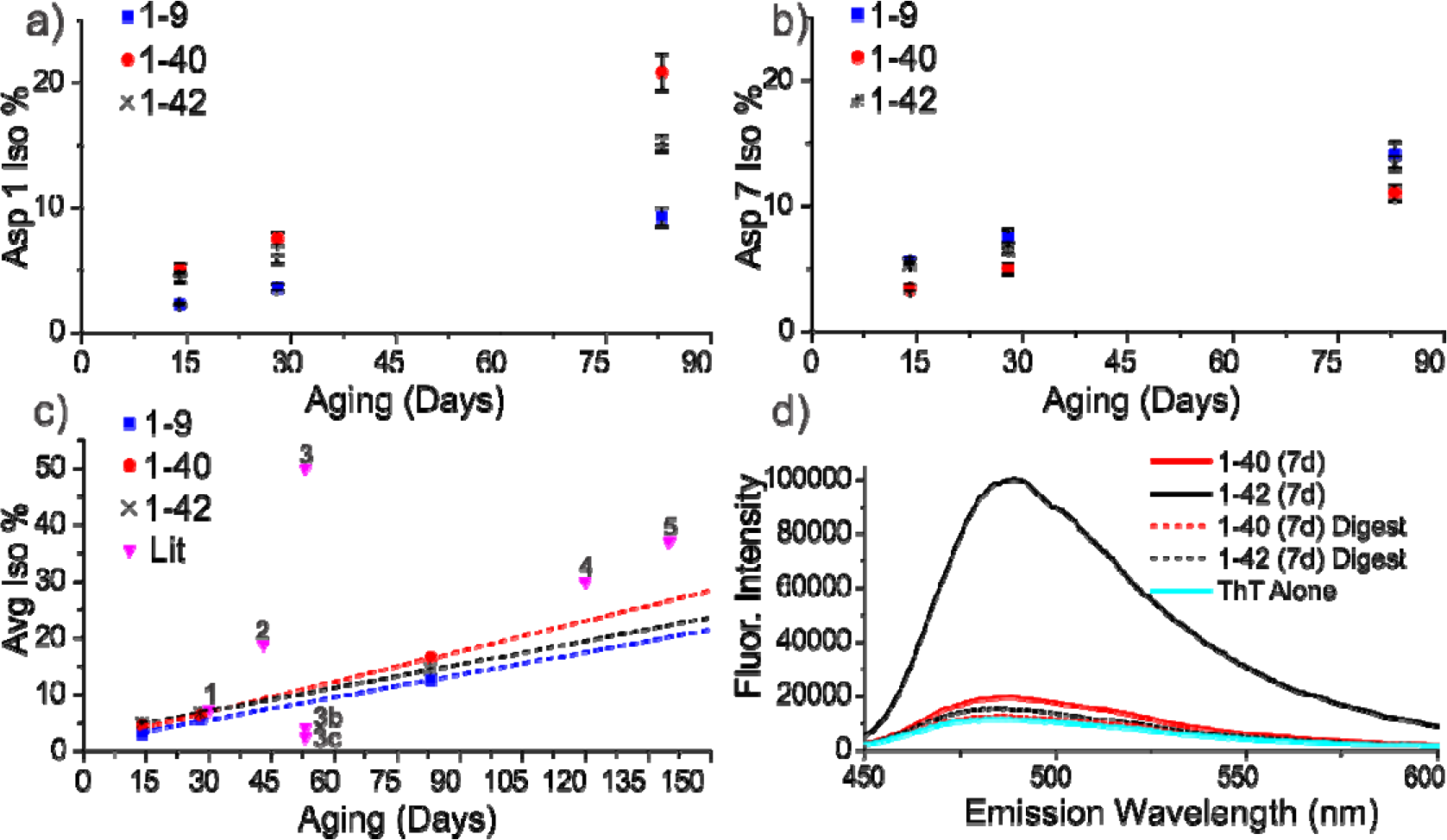
Isomerization % as function of time for a) Asp1 and b) Asp7. c) Average isomerization rate for Asp1 and Asp7 relative to rates from literature. d) ThT assay after 7 days confirming that any fibrils are largely digested during analysis.

These experiments were conducted at µM concentrations, which is sufficient for the formation of amyloid fibrils. The presence of amyloid was examined by ThT assay after 7 days as shown in Fig. 4d. The assay reveals that Aβ1-42 had already formed fibrils within 7 days, while Aβ1-40 was just entering fibril formation, consistent with previous reports.^46^ After digestion with chymotrypsin, the fluorescence diminishes substantially, suggesting that fibrils are broken up and should not significantly influence the analysis. Interestingly, amyloid formation appears to slightly increase the rate of isomerization for Asp1 but in general does not significantly influence the rates.

### A framework connecting lysosomal failure and AD

Long-lived proteins are subject to many spontaneous chemical modifications, including subtle changes such as iso/epi modifications that may seem harmless and are easily overlooked. Nevertheless, heavy isotope pulse-chase experiments in mice have shown that long-lived proteins in the brain are more commonplace than previously realized and can persist for timespans exceeding one year.^47^ These long-lived proteins are part of the overall equation that must be balanced to maintain proteostasis and will therefore be targeted for degradation at some point. Our results reveal that isomerized and epimerized sites in long-lived proteins resist digestion by the primary cathepsins present in lysosomes. Both epimerization (Ser and Asp) and isomerization (Asp) effectively prevent proteolysis at the site of modification and nearby residues for both endo- and exopeptidases. Long-lived proteins targeted to the lysosome are therefore expected to produce residual peptide fragments that are too long to be recognized by the transporters responsible for releasing digested amino acids back to the cytosol. Additionally, the residual peptides will contain an unnatural amino acid that would be expected to further frustrate transporter recognition. Accumulation of these byproducts within the lysosomal machinery is therefore possible. In fact, interference with lysosomal function has already been documented in similar circumstances with pyroglutamate modified Aβ, where the influence on proteolysis is significantly less pronounced.^48^

We have demonstrated that iso/epi modifications significantly inhibit lysosomal digestion in glial cells, but prior work has additionally shown that such modifications are toxic. Makarov and coworkers have examined isomerization of the N-terminal portion of Aβ in relation to the idea that such modifications enhance amyloid formation in the presence of zinc ions. They found that isomerized Aβ1-42 was more toxic than the canonical form when incubated with several different cell lines (NSC-hTERT, SK-N-SH, and SH-SY5Y).^49^ Furthermore, cell death by apoptosis rather than necrosis was more prevalent in the case of isomerized Aβ, indicating an alternate and more specific mechanistic pathway. Importantly, related experiments have demonstrated that Aβ localizes into the lysosome when incubated with SH-SY5Y cells,^50^ suggesting that the toxicity could be reasonably attributed to lysosomal pathology instead. Toxic effects have also been found in animal studies.^51^ Perhaps most strikingly, injection of isomerized Aβ1-16 leads to significantly increased amyloid plaque accumulation in 5XFAD transgenic mice whereas canonical Aβ1-16 does not.^52^ Importantly, Aβ1-16 does not contain the amyloid forming portion of the peptide. Although these data could be interpreted to support to the zinc-mediated amyloid aggregation hypothesis, our findings suggest that disruption of the lysosomal system could also explain the results. Introduction of isomerized Aβ1-16 could lead to lysosomal failure, followed by disrupted proteostasis and the observed increase in amyloid plaque formation.

We have established that isomerization of Aβ is relatively fast. The residence time of Aβ in the human brain is difficult to determine due to the multiple destinations and pathways that can be taken, but studies have shown that the fraction of Aβ escaping into cerebrospinal fluid persists beyond 30 hours in a healthy individual.^53^ Similar studies have shown that clearance rates for Aβ are mismatched relative to production in AD individuals,^54^ which suggests that some fraction evades degradation and may persist for longer times. The rates in Fig. 4 allow for a small degree of isomerization (~0.2%) even within a 30 hour timeframe. Furthermore, any fraction of Aβ residing in the brain for a week or more would be expected to isomerize significantly. The N-terminal region of Aβ is disordered in amyloid structures determined by NMR,^55,56^ which may allow free access to the required succinimide intermediate while providing some catalytic interactions that favor isomerization. Aβ is therefore a likely source of isomerized residues in the brain, but a few reports have shown that tau can also be isomerized due to deamidation at positions 596 and 698, or isomerization of Asp at positions 510 and 704.^57,58^ The size and largely unstructured nature of tau^59^ make it almost certain that other sites of isomerization also exist. There is ample evidence that the proteins most strongly associated with AD pathology are subject to iso/epi modifications that could lead to lysosomal failure.

## Conclusion

Iso/epi modifications are clearly generated on a relatively short timescale and prevent cathepsin digestion of nearby peptide bonds. Although other proteolytic pathways exist within cells that may also encounter difficulties with iso/epi modifications, lysosomes are uniquely vulnerable because undigested byproducts cannot escape the lysosomal membrane and can eventually cause failure and storage of the entire organelle. When this sequence of events is triggered in lysosomal storage disorders, the consequences are dramatic and often fatal. Malfunction of the lysosome is also strongly associated with the pathology of AD, as are misfolding and aggregation of both Aβ and Tau. Lysosomal failure caused by the iso/epi modifications documented to exist in both Aβ and Tau offers a direct connection between these observations and a potential new pathway to explore for the underlying cause and treatment of AD.

## Acknowledgements

The authors are grateful for funding from the NIH (R01GM107099 to RRJ, and R01NS091616, R21NS106949, R25GM119975 to BDF). Min Xue is kindly thanked for allowing us to use his fluorescent plate reader. Hill Harman, Gal Bitan, Pablo Martinez, and Joe Loo are acknowledged for helpful discussions.

## Supporting Information

### Materials and Methods

Scheme S1 depicting the spontaneous deamidation and isomerization pathway. Fig S1 additional digestion rate data.

Fig S2 LCMS data comparing chromophoric and native peptide digestion efficiency.

