## supporting info for "Spontaneous isomerization of long-lived proteins provides a molecular mechanism for the lysosomal failure observed in Alzheimer’s disease" (from another version of this preprint: version 1, 2019-04-12)

**Materials and Methods**

Peptide Synthesis

Peptides were synthesized following an accelerated FMOC-protected solid-phase peptide synthesis protocol.^[1]^ Fluorophores Edans and Dabcyl were purchased as FMOC-protected amino acid conjugates and attached to peptides during synthesis. Following synthesis, peptides were purified on a Varian ProStar HPLC using a Phenomenex Jupiter Proteo C12 4 μm 90 Å 250 mm × 4.6 mm column. Purified peptides were stored frozen in 50/50 acetonitrile/water (v/v) and vacuum-dried before reconstitution in appropriate buffers.

In Vitro Aging

Lyophilized amyloid beta powder was disentangled via ammonium hydroxide treatment.^[2]^ Following disentanglement, aliquots containing 100 μg amyloid beta were lyophilized before reconstitution in 1 mL 50 mM Tris 7.4 pH at 37 °C. Samples were aged in microcentrifuge tubes in a sand-bath style incubator. Aliquots were removed at given timepoints and frozen until analysis. Aged samples were digested at room temperature for 18 h with chymotrypsin in 200 mM Tris buffer with 4 mM CaCl_2_ and an enzyme:substrate weight ratio of 1:2 to ensure complete digestion of fibrils. Digested samples were frozen until analysis.

LCMS Analysis

Samples were analyzed on an Agilent 1100 HPLC interfaced with a Thermo LTQ or Thermo Fisher Velos Pro Orbitrap using electrospray ionization sources. Peptides were separated on a Thermo Scientific BetaBasic C-18 3 μm, 150 x 2.1 mm, column. Mobile phase A was water with 0.1% formic acid, and mobile phase B was methanol with 0.1% formic acid.

In Vitro Fluorimetry

Aliquots containing 75 μg of purified peptide were added to a 96-well plate in 50 mM acetate buffer pH 5.5 with 1 mM EDTA and 1 mM DTT to prevent active site oxidation. Cathepsin L was added at an enzyme:substrate ratio of 1:150. Digestion was then monitored using a Biotek Synergy H1 fluorescent plate reader with an excitation wavelength of 336 nm and emission wavelength of 490 nm. Fluorescence is plotted as the average of 3 runs and error bars represent the standard deviation.

The presence of amyloid fibrils was examined by ThT assay.^[3]^ Amyloid beta samples were diluted to 2 μM in Tris buffer before addition of 30 μM ThT. Emission scans were performed on a QuantaMaster-400 fluorimeter, using an excitation wavelength of 440 nm.

In Vitro Digestion

Aliquots containing 75 μg of purified peptide were dissolved in 150 μL of buffer. Digestions utilizing cysteine cathepsins (CatL, CatB, CatH) were performed in 50 mM acetate buffer pH 5.5 with 1 mM EDTA and 1 mM DTT to prevent active site oxidation. Digestions with CatD were performed in 50 mM acetate buffer pH 4.5 to ensure protonation of the active site aspartic acid. Enzyme ratios were adjusted based on activity; CatL and CatB were used at an enzyme:substrate ratio of 1:150, CatD 1:50, and CatH 1:25. Timepoint aliquots were taken at 0.5, 2, 4, 6, 8, 24, and 48 hours, and diluted to a final concentration of 5 μM with 0.2% TFA solution to pH quench the enzymatic reaction. Digested samples were frozen until analysis.

Microglial Cell Culture and Imaging

SIM-A9 microglial cells were seeded into 8-well chamber slides (Thermo Scientific, 154941) at a density of 2x 10^5^ cells/cm^2^ in complete media (DMEM:F12 + 10% FBS + 5% Horse serum + 1% penn/strep) and allowed to attach for 12 hours under standard conditions (37°C, 5% CO_2_). At the start of experiments, 10 µM of either L- or D-isoAsp peptide was added. Digestion of the cell-penetrating peptide isomers was monitored in live cells on a Zeiss 880 laser scanning confocal microscope. The sample chamber was maintained at 37 °C for the duration of the experiments. Images were collected using a Zeiss Plan-Apochromat 10x/0.45 M27 objective. Eight 850 μm x 850 μm images were collected and tiled as two sets of 2x2 images. Fluorescence quantitation is derived from the full image. The data presented in Fig 3b encompasses 936 cells for the L-Asp peptide incubation (average = 234 per image, stdev = 36) and 1282 cells for the D-isoAsp peptide incubation (average = 320 per image, stdev = 51). Data from a second location Fig. S1c which was also collected as 2x2 tiled images, and encompasses 986 cells for the L-Asp peptide incubation (average = 247 per image, stdev = 26) and 695 cells for the D-isoAsp peptide incubation (average = 174 per image, stdev = 20). Fluorescent signal was plotted as fluorescence per cell after establishing a signal threshold based on the negative control. Brightfield images were collected for cell-counting, and digestion was monitored by recording Edans derived fluorescent emission from 481-499 nm. Following imaging, cells were counted in FIJI,^[4]^ using a threshold image approach that was validated manually. Raw fluorescence data was recorded for both peptide isomers in addition to a negative control (ie cells without peptide). To eliminate background noise, scattering, and autofluorescence, only fluorescent signal that is above the negative control is used for quantitation.

Molecular Modeling

The active site of cathepsin L was modeled in Maestro 11 (Schrödinger, LLC). The PDB deposit, 3K24, was processed to include H-bonds and displayed using the surface representation. The QLA peptide bound in the crystal structure was modified to QLD and evaluated for potential H-bonds as well as steric clashes. The chirality of the alpha-carbon of the Asp residue was inverted to generate the D-Asp isomer.

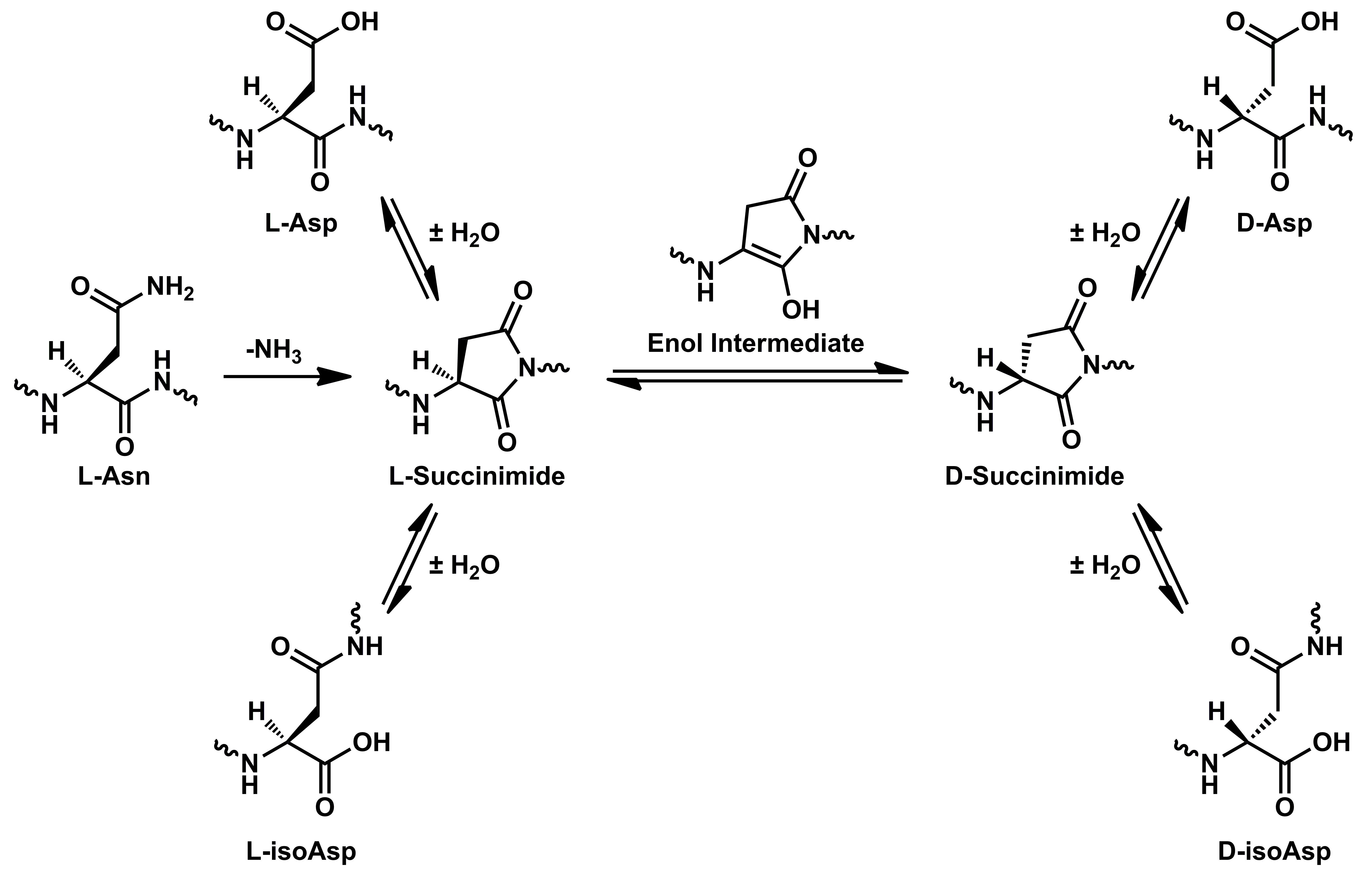

**Scheme S1**. Pathways for isomerization of aspartic acid and deamidation of asparagine.

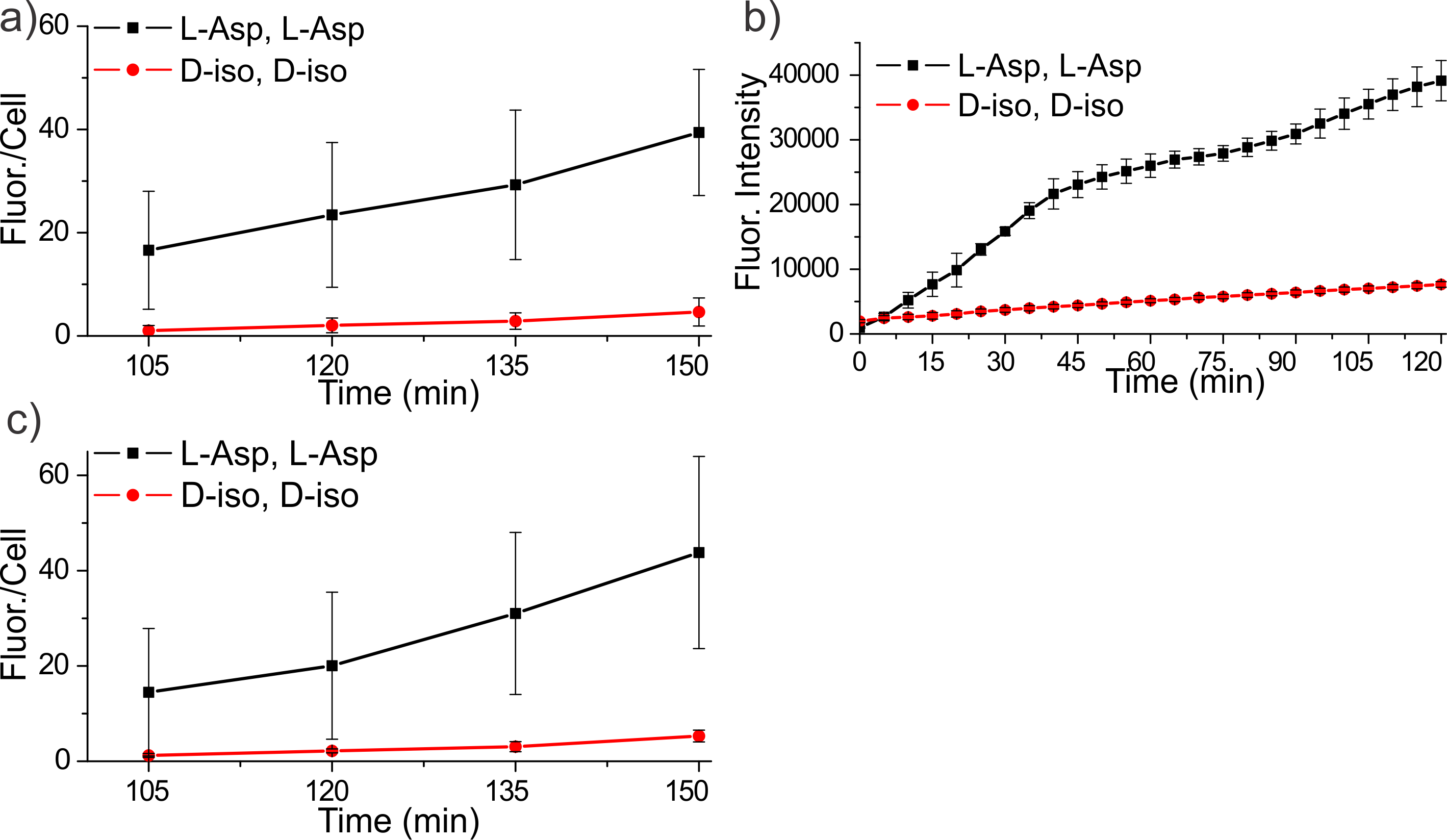

**Figure S1.** Comparison of the microglial digestion rate data from Fig 3b replotted in the same format as the data shown in Fig 3c. a) replotted data, b) original Fig 3c for easy comparison. Values represent averages with error bars showing standard deviations. c) Similar results obtained from a second, independent location.

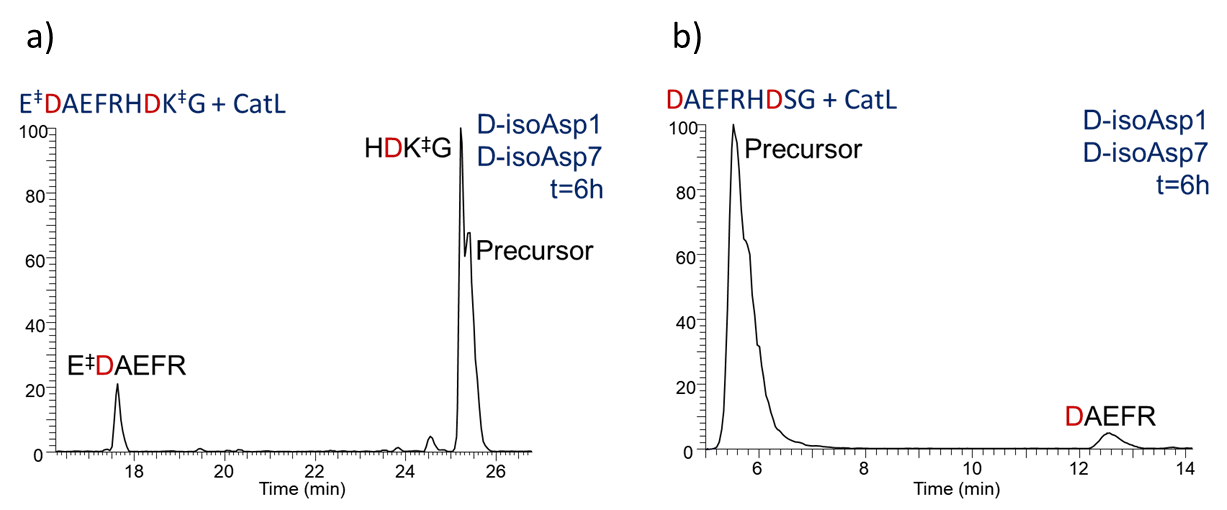

**Figure S2.** Comparison of digestion of a) chromophoric and b) native-state peptides by LC-MS. Digestion proceeds more rapidly after addition of chromophoric probes for cell imaging.
